# Exploring the Human-Coronavirus protein-protein interaction network from the perspective of a novel host-virus association

**DOI:** 10.1101/2024.01.12.575398

**Authors:** Debarun Acharya, Tapan K Dutta

## Abstract

Host-pathogen interaction is the best example of an evolutionary arms race where pathogen and host continuously coevolve to survive and exert negative effects on each other. The adaptability of both host and pathogen is critical for this association. In this study, we explored the association of severe acute respiratory syndrome (SARS) coronaviruses (CoVs) with their human host from the genomic and evolutionary perspectives based on a comparative analysis of SARS and MERS coronaviruses. We observed that human proteins that are part of the SARS-CoV2-human association are enriched in hubs and bottlenecks. Again, these proteins take part in more protein complexes and show slower evolutionary rates compared to the human proteins associated with the two other coronaviruses, SARS-CoV and MERS-CoV. Moreover, the human proteins involved in the interaction with SARS-CoV2 are mostly longer proteins harboring long intrinsically disordered stretches and a higher level of disordered protein binding sites. Codon usage analysis revealed that the novel coronavirus is least adapted to codons used in housekeeping and lung-specific genes, compared to the other two coronaviruses. We conclude that the signatures showed by the SARS-CoV2-human protein interaction network revealed the virus’s association with vital human proteins and pathways, via interactions mediated by protein complexes and intrinsically disordered protein binding sites, which may have assisted the higher infectivity of SARS-CoV2 in its human host than the other two less-virulent human coronaviruses, despite having a lower optimization to its host’s codons.

## 1. Introduction

The recent COVID-19 pandemic caused by the novel coronavirus SARS-CoV2 has been a serious threat to global public health, with its effects persisting throughout most countries and continents. Since the first occurrence of the SARS-CoV2 infection in 2019, a plethora of research articles have been published to explore the origin and the nature of the virus (Hu et al., 2021; Li et al., 2020; Pagani et al., 2023; Rasmussen, 2021; Singh & Yi, 2021), the molecular basis underlying the pathogenesis and infection (Jackson et al., 2022; Lamers & Haagmans, 2022; McCallum et al., 2021; Niu et al., 2021; Shang et al., 2020), the epidemiology of the disease (Alteri et al., 2021; Dhar et al., 2021; Esper et al., 2021), the impact on human population (Aburto et al., 2021; Aburto et al., 2022; Jin et al., 2021), the protection strategies from the virus (Abdoul-Azize & El Gamil, 2021; Overbaugh, 2020; Zheng et al., 2021), and the genetic variations and evolution (Li et al., 2020; Markov et al., 2023; Roemer et al., 2023) among the many aspects of SARS-CoV2 infection. The high infectivity and mortality rates among SARS-CoV2 patients influenced these studies and accelerated the development of therapeutic and prophylactic measurements against SARS-CoV2 throughout the globe (Andreano & Rappuoli, 2021; Hotez et al., 2021). This pandemic situation further initiated the discovery of efficient treatments like convalescent plasma therapy (Chen et al., 2020; Focosi et al., 2020), the use of monoclonal antibodies (Lloyd et al., 2021; Marovich et al., 2020; Wang et al., 2021), the development of vaccines (Krammer, 2020; Ndwandwe & Wiysonge, 2021; Tregoning et al., 2020), and the repurposing of drugs (Consortium, 2021; Jang et al., 2021; Li & De Clercq, 2020; Tarighi et al., 2021; Wang et al., 2020; Y. Zhou et al., 2020). About 300 repurposed drugs (Menestrina et al., 2021) and 50 vaccines (https://covid19.trackvaccines.org/vaccines/approved/#vaccine-list) were approved globally to combat the deadly virus as of June 2025. However, the continuous accumulation of mutations in SARS-CoV-2 has posed significant challenges to therapeutic strategies, contributing to the emergence of multiple variants— from Alpha and Beta in December 2020 to Omicron in November 2021 and more recently to subvariants such as JN.1, KP.3, KP.3.1.1, LB.1, XEC, LP.8.1, and NB.1.8.1 observed throughout 2023–2025—many of which exhibit increased transmissibility or immune evasion potential, showing a high adaptability of this virus (Lippi et al., 2022; Roemer et al., 2023; Singh et al., 2021; Who, 2025).

The horseshoe bats (*Rhinolophus* spp.) are the reservoir hosts of coronaviruses, including the SARS-CoV and SARS-CoV2 (Ruiz-Aravena et al., 2021; Wang et al., 2006; P. Zhou et al., 2020), with a few reports indicating pangolins as the intermediate host (Lam et al., 2020; Liu et al., 2020; Zhang & Holmes, 2020). The high genomic sequence identity (∼99.99%) of SARS-CoV2 viruses isolated from different patients suggests a very recent host shift to humans. Although both SARS-CoV and SARS-CoV2 share their entry receptor, angiotensin-converting enzyme 2 (ACE2), that binds to the spike protein of both coronaviruses, the affinity for ACE2 is 10-20 times higher for the SARS-CoV2 spike protein than the SARS-CoV counterpart explaining the higher infectivity of the former (Abdolmaleki et al., 2022).

The protein-protein interactions are crucial components in the molecular crosstalk between host and pathogen during infection. Such interactions are necessary for the establishment of the pathogen and its multiplication inside the host. For a novel pathogen, interactions are often random and may lead to severe consequences in the host. However, from the perspective of pathogens, killing the host is suicidal, and extreme virulence is, therefore, not a sustainable evolutionary solution (Fieldman, 2024). Instead, the pathogen undergoes evolutionary selection within the host’s body, gradually adapting to the host environment over time. This process shapes the coevolutionary dynamics of host-pathogen interactions. Thus, the increasing multiplication and exposure to the human subpopulations provides the novel virus a large evolutionary space to form mutant variants, evolutionarily equipped to survive and reproduce (Lippi et al., 2022; Singh et al., 2021).

The novel coronavirus SARS-CoV2 is the newest member of the *Coronaviridae* family, belonging to the *Sarbecovirus* subgenus (genus *Betacoronavirus*) (Lefkowitz et al., 2018). It is a close relative to the animal coronaviruses discovered early in the twentieth century and the third coronavirus known to be pathogenic for humans (Sironi et al., 2020). Although, until recent times the effect of human-infecting coronaviruses (namely Human coronaviruses HCoV-229E, HCoV-OC43, HCoV-NL63, and HCoV-HKU1) was only limited to mild symptoms comparable to the common cold (Liu et al., 2021). On the other hand, the outbreak of severe acute respiratory syndrome (SARS) coronavirus in 2003 (SARS-CoV) caused the death of more than 700 people globally. The Middle East Respiratory Syndrome Coronavirus (MERS-CoV) was initially identified in Saudi Arabia in 2012. This virus follows a zoonotic mode of transmission, being spread to humans from dromedary camels that were their major reservoir host (Omrani et al., 2015). The disease has a high death rate of about 35% of the infected persons, affecting about 1400 individuals in South Korea and Middle East countries, like Saudi Arabia, United Arab Emirates, Jordan, Qatar, Oman, and Iran. The severity of infections caused by these viruses is much lower compared to SARS-CoV2, as the recent pandemic caused by SARS-CoV2 led to over 7 million deaths and about 780 million infected persons globally as of June 2025. The SARS-CoV2 has 79.7% sequence similarity with the SARS-CoV of 2003 (Y. Zhou et al., 2020), the highest compared to other pathogenic coronaviruses. The devastations were as extreme as more than 100,000 global deaths per day. Apart from the huge number of casualties, the socioeconomic distress caused by the pandemic is likely to last longer than anticipated.

Additionally, it does not change the fact that the interactions with human proteins for both the SARS-CoVs and MERS-CoV may control the severity of these infections, influencing the biological function of the human proteins and their associated pathways, via the protein-protein interaction (PPI) network. Moreover, other preventive measures of the disease such as passive immunization or vaccination certainly help to stop the disease transmission or may imply further selection pressure to generate vaccine-resistant variants (Luo et al., 2021). The underlying cause of rapid propagation of SARS-CoV2 across human populations suggests that SARS-CoV2 may have an upper hand in regulating the human PPI network, via the establishment of a stable pathogen-host PPI network. Comprehensive host-pathogen protein-protein interaction network analysis for human pathogens have suggested that pathogen proteins often interact with human proteins with a high degree centrality, which is consistent for viral, bacterial, protozoan, and fungal pathogens (Acharya & Dutta, 2021; Crua Asensio et al., 2017; Dyer et al., 2008; Nithya et al., 2021). Considering the coronaviruses, the novel coronavirus SARS-CoV2 has recently shown a global outbreak, and a more rapid adaptation than the other two coronaviruses (MERS-CoV and SARS-CoV) in a very short span. As a result, this rapid adaptation of the SARS-CoV2 should be evident in the pathogen-human protein-protein interaction (PHPPI) network involving the human and SARS-CoV2 proteins. Using high-throughput affinity purification mass spectrometry, Gordon et al. first mapped the interactions between 26 SARS-CoV2 proteins and 332 human proteins, revealing key host targets involved in translation, vesicle trafficking, and immune signaling (Gordon et al., 2020). Such analyses help identify functional protein modules and prioritize druggable host factors, offering potential therapeutic targets beyond viral components. Integration of SARS-CoV-2 PPIs with host systems biology—such as protein complexes and signaling pathways—further enables understanding of the broader cellular impact and network perturbations caused by infection (Stukalov et al., 2021). On the contrary, the MERS-CoV has mostly circulated among humans from camels and SARS-CoV circulated among humans for a very short time (Omrani et al., 2015) and their association with human proteins in the PHPPI network should reflect a weaker association. Concerning the higher severity of SARS-CoV2 infections, SARS-CoV2 may have a higher control over host cellular systems, despite being novel, due to its rapid adaptability and propagation within human populations. In this study, we have analyzed the recent human coronavirus interactome data for MERS-CoV, SARS-CoV and SARS-CoV2 from the perspective of protein-protein interactions between human and coronaviruses. The study aims to compare these three human coronaviruses from a network biology perspective accounting for the roles of human proteins, their network topology and structural features that facilitate these interspecies interactions.

## 2. Materials and methods

### 2.1. Sequence data

The coding sequences (CDS) for MERS and SARS coronaviruses were obtained using the NCBI GenBank (Benson et al., 2012) (NCBI Reference Sequences NC_019843.3, NC_004718.3 and NC_045512 for MERS-CoV, SARS-CoV and SARS-CoV2, respectively). The Human CDS sequences were obtained using the Biomart interface of Ensembl (Yates et al., 2020). Human protein sequences were obtained from the UniProt (UniProt, 2015) [proteome ID: UP000005640]. We have selected the ‘reviewed’ proteome only, which leads to a list of 20410 proteins and their corresponding amino acid sequences.

### 2.2. Protein-protein interaction data

The coronavirus-human protein-protein interaction data for MERS-CoV, SARS-CoV, and SARS-CoV2 were obtained from H2V (Zhou et al., 2021) and BioGRID (Oughtred et al., 2021) database. In BioGRID, we have filtered the interaction data and included only experimentally validated PPI data involving large-scale experiments (involving >10 PPI pairs) that show direct evidence of interactions, by incorporating the following experimental methods: Affinity Capture-MS, Affinity Capture-Western, Biochemical Activity, Co-crystal Structure, Cross-Linking-MS, PCA, Proximity Label-MS, Reconstituted Complex and Two-hybrid. We have integratred the filtered human-virus interacting pairs obtained from H2V and BioGRID databases and removed the redundant interacting pairs. The final list of Human-coronavirus PPI data contains 694 Human-MERS-CoV, 1451 Human-SARS-CoV and 12134 Human-SARS-CoV2 binary protein-protein interaction pairs, involving 572, 1142 and 3168 human proteins, respectively. The proteins interacting with both SARS-CoV and SARS-CoV2 (N= 490) were also considered, but the proteins interacting with all coronaviruses (MERS-CoV, SARS-CoV and SARS-CoV2), and those interacting with MERS-CoV and any SARS coronavirus (MERS-CoV and SARS-CoV, MERS-CoV and SARS-CoV2) were discarded to remove any influence of MERS-CoV in the results. The proteins without any involvement of interaction of any of the coronaviruses under study were considered as a ‘Noninteracting’ group (N= 16566).

The human protein-protein interaction network was obtained from the supplementary data of Cheng *et al*. (Cheng et al., 2018), which provides a comprehensive list of human protein-protein interaction networks collected from established PPI databases. With this data, we have included the updated human binary protein-protein interaction data from BioGRID (version 4.4.245) (Oughtred et al., 2021), which includes only the validated PPIs from experiments showing direct evidence of interaction, as mentioned earlier. The Entrez gene IDs were converted to corresponding UniProt IDs using UniProt’s ID conversion tool (UniProt, 2015). The final data contains 372895 interactions involving 16358 human proteins. The human PPI network was analyzed using the NetworkAnalyzer plugin of Cytoscape 3.10.1 (Shannon et al., 2003) to determine the degree and betweenness centrality of human proteins. The top 20% proteins having the highest degree and betweenness centrality were considered hubs and bottlenecks, respectively (Acharya & Dutta, 2021).

### 2.3. Protein complex association

The human protein complexes were obtained from the Corum database (release 5.1) (Tsitsiridis et al., 2023). The human protein complex data provides a unique number for each protein complex and the UniProt ID of the participants of the complex. The unique number of complexes involving a protein was considered as its complex number, which was calculated using in-house Python script. We have obtained a list of 5366 human protein complexes involving 5051 human proteins that have matched with 4799 proteins in total from the five groups of our study.

### 2.4. Protein intrinsic disorder and disordered protein binding sites

The intrinsic disorder of human proteins was predicted using the IUPred2A (Mészáros et al., 2018), where each amino acid residue in a protein is provided with a disorder score ranging from 0 to 1. In IUPred, the residues with a score of ≥ 0.5 are considered disordered residues, and the consecutive stretches of ≥ 30 amino acid-long disordered residues are considered long-disordered regions. An in-house PERL script was used to determine such long-disordered regions of human proteins, with a consecutive stretch of 30 or more amino acids having a disorder score of ≥ 0.5 for all the amino acids in the stretch (Acharya & Dutta, 2021). The long-disordered protein binding regions were calculated using the ANCHOR2 under IUPred2A, an additional tool designed to predict disordered binding regions within intrinsically disordered regions. For each protein sequence, ANCHOR2 provides a per-residue score indicating the likelihood of that residue participating in a disordered protein-protein interaction interface. Disordered stretches within a protein containing short stretches of amino acid residues with ANCHOR2 scores ≥ 0.5 were classified as the putative disordered protein binding regions. Two quantitative metrics were extracted from each protein’s ANCHOR2 profile: the ANCHOR regions: the total number of regions with ≥ 5 residues with ANCHOR2 scores ≥ 0.5 within the disordered stretches and total ANCHOR residues per proteins that fall within the disordered residues and have ANCHOR2 scores ≥ 0.5 (Dosztányi et al., 2009). In addition, the Molecular Recognition Features (MoRFs) of human disordered proteins were obtained from the fMoRFpred webserver (Yan et al., 2016), where each amino acid in a disordered protein is classified as MoRF or non-MoRF residue. Similar to ANCHOR, the consecutive stretches of ≥ 5 MoRF residues were considered as regions responsible for disordered protein binding (MoRF regions) and the number of such regions in each human protein, along with the total MoRF residues in a protein was calculated using an in-house PERL script.

### 2.5. Evolutionary Rate

The evolutionary rates of human proteins were obtained using the Ensemblbiomart (Yates et al., 2020), using one-to-one orthologs of human proteins with four other species: Chimpanzee (*Pan troglodytes*), Macaque (*Macacamulatta*), Pig (*Susscrofa*), and Mouse (*Musmusculus*). We obtained the nonsynonymous nucleotide substitutions per nonsynonymous sites (dN) and synonymous nucleotide substitutions per synonymous sites (dS) of the orthologous pairs from Ensembl biomart. The dN/dS ratio for each human protein, with their 1:1 orthologous proteins in all the species considered, were used to measure the evolutionary rate. The mutation saturation was controlled by removing all proteins having dS values >3 (Acharya & Ghosh, 2016).

### 2.6. Pathway enrichment analysis

The pathway enrichment analyses were conducted using the clusterProfiler R package(Yu et al., 2012). The UniProt IDs were converted to Entrez Gene IDs using the org.Hs.eg.db annotation database in R (Carlson et al., 2019). The resulting gene sets were grouped based on viral interaction status (MERS-CoV, SARS-CoV and SARS-CoV2), producing three distinct input lists for comparative analysis. The pathway enrichment analysis was conducted using Gene Ontology (GO) enrichment for the Biological Process (BP) ontology (Gene Ontology, 2019) using the enrichGO function, with the Benjamini–Hochberg method for multiple testing correction. We have also performed KEGG Pathway (Kanehisa et al., 2025) enrichment using the enrichKEGG function and adjusted p-values via the Benjamini–Hochberg method. Enrichment results for both GO terms and KEGG pathways were visualized using a dotplot from the enrichplot package. Dot plots were faceted by the virus group to enable direct visual comparison of the top ten significantly enriched terms.

### 2.7. Housekeeping and lung-specific genes

Human housekeeping genes were obtained from the HRTAtlas 1.0 database (Hounkpe et al., 2021), which presents the human housekeeping genes that are stably expressed in 52 tissues and cell types. We obtained a final list of 2071 non-redundant protein-coding housekeeping genes. The genes expressed in the lungs were obtained from the Human Protein Atlas using the Tissue Atlas for lung transcriptome (Uhlén et al., 2015). We have selected a final list of 190 protein-coding genes that are selectively elevated in the lungs.

### 2.8. Codon usage

The frequency of each codon was calculated using the ‘cusp’ function of the EMBOSS, an open-source software package (Rice et al., 2000). The function ‘cusp’ calculates the codon usage for one or more nucleotide coding sequences and provides a tabulated output, using desired CDS sequences. The codon composition of human coronavirus CDS and that of the human housekeeping and lung-specific genes were calculated using the ‘cusp’ function. Similarly, the CDS sequences of viral proteins for the three coronaviruses were used to determine their codon usages. The expected number of each of the 61 codons in the provided input sequence, per 1000 bases (i.e., codon frequency) was correlated for each coronavirus CDSs with that of human housekeeping (HK) and lung-specific (Lung) genes, thereby obtaining the Pearson correlation coefficient (PCC) for each combination (HK-MERS-CoV, HK-SARS-CoV, HK-SARS-CoV2 and Lung-MERS-CoV, Lung-SARS-CoV, Lung-SARS-CoV2).

### 2.9. Statistical Packages and Software

The statistical analyses have been carried out using IBM SPSS Statistics 22 (Meyers et al., 2013), R packages. Statistical analyses were conducted using non-parametric methods due to the non-normal distribution of the data. For comparisons between two independent groups, the Mann-Whitney U test was applied to evaluate differences in central tendency. For comparisons involving more than two independent groups, the Kruskal-Wallis H test was used to assess whether group medians differed significantly. Jonckheere-Terpstra test was conducted to detect ordered differences across groups. All statistical tests were two-tailed, and a p-value of less than 0.05 was considered indicative of statistical significance. The codon usage correlation analyses have been carried out using the Pearson correlation coefficient. In house PERL, Python and R scripts were also used in data analyses.

## 3. Results

### 3.1. Network centrality of human proteins interacting with coronaviruses

Human proteins were differentiated based on their interaction statuses with the proteins of human coronaviruses, resulting in 2302 human proteins interacting with SARS-CoV2 only, 480 proteins interacting with SARS-CoV only, 490 proteins interacting with both the SARS-CoVs, 146 proteins interacting with MERS-CoV only, and 16566 non-interacting proteins which imply no reported interaction with any of these coronaviruses (see materials and methods). The central proteins in the human protein-protein interaction (PPI) network were identified from the constructed human high-throughput PPI data, from where the hubs and the bottlenecks were classified using degree and betweenness centrality values, respectively (see materials and methods). Our comparative analysis revealed that the human proteins that do not interact with any coronavirus proteins (noninteracting group) contain the lowest proportion of hubs and bottlenecks (Figure 1). Among the coronavirus-interacting groups, human proteins interacting with SARS-CoV2 contain a higher proportion of hubs and bottlenecks, followed by MERS-CoV and SARS-CoV. The result indicates two observations: firstly, the human coronaviruses mostly target host central proteins with higher connectivity, as revealed in earlier studies with other pathogens (Calderwood et al., 2007; Dyer et al., 2008). Secondly, the SARS-CoV2-interacting human proteins have a higher proportion of hubs and bottlenecks than those interacting with other two coronaviruses. This suggests that the SARS-CoV2 may have a better opportunity to hijack the host cellular network more efficiently compared to the other two coronaviruses under study, which may be helpful for the optimization of viral propagation and virulence, despite its recent zoonotic origin.

**Figure 1.**
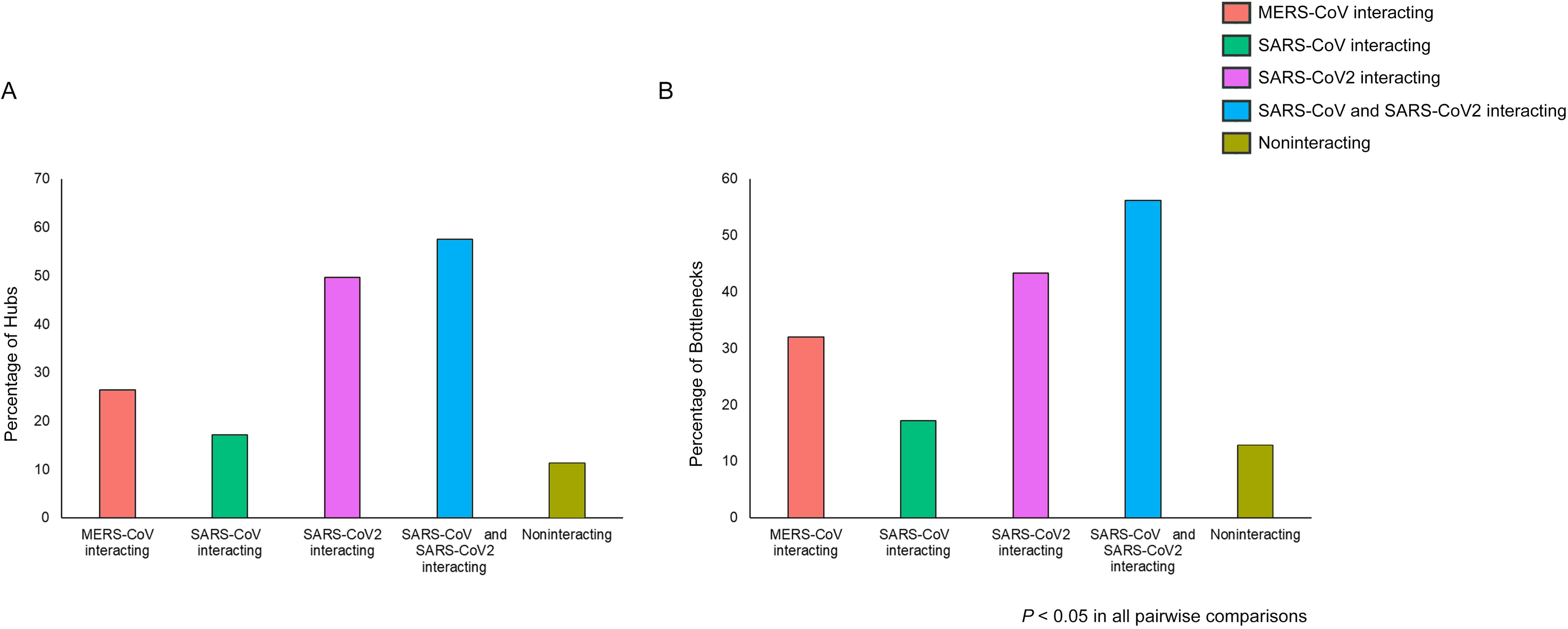
Percentage of **A.** hubs and **B.** bottlenecks in human proteins, grouped according to the coronavirus interacting status. All the differences are statistically significant (*P* < 0.05).

### 3.2. Protein complex association

The human cellular interaction network is composed of many proteins that often form complexes that perform particular cellular functions with the help of its participants. The protein complex association reflects how a protein is integrated into various macromolecular complexes to carry out the specific functions of those complexes. The protein’s participation in multiple complexes, referred to as the protein complex number (PCN) (Chakraborty & Ghosh, 2013), serves as an appropriate measure of the number of functions (multifunctionality) in which the protein is involved. Proteins that are involved in many complexes, are important molecular players of the cell. Here, the protein complex association of human proteins interacting with different coronaviruses revealed that the complex number of human proteins follows the following trend regarding the five groups under study: SARS-CoV2 (PCN= 4.149) > SARS-CoV_and_SARS-CoV2 (PCN= 3.836) > MERS-CoV (PCN= 3.658) > SARS-CoV (PCN= 3.341) > Noninteracting (PCN= 3.363). To assess the significance of this directional trend in protein complex number across the ordered viral interaction groups, the Jonckheere-Terpstra test was employed to confirm a statistically significant increasing trend (*P*= 1.43 × 10^−2^, Jonckheere-Terpstra test). This suggests that the virus-interacting human proteins have a higher protein complex association. Moreover, among these viruses, SARS-CoV2 proteins can hijack proteins participating in more complexes within the human PPI network. Nevertheless, to survive within the human host, the interactions between these host proteins and viral proteins are crucial, which warrants the exploration of the structural features of host proteins that are assisting in such interactions and the relevance of such features considering human-coronavirus interactions.

### 3.3. Intrinsic disorder and disordered protein binding regions

Protein intrinsic disorder plays an important role in the host-pathogen protein-protein interaction network as it provides flexibility to protein structure to promote low-affinity interactions between proteins (Acharya & Dutta, 2021; Panda et al., 2017). The increased flexibility of the host’s disordered proteins allows host-pathogen protein interactions, which otherwise would remain infeasible (Acharya & Dutta, 2021). For a detailed understanding of this scenario, human-coronavirus protein-protein interactions were examined using the IUPred2A (Mészáros et al., 2018) protein intrinsic disorder prediction algorithm with regards to the coronaviruses interacting and noninteracting groups. Interestingly, the human proteins interacting with SARS-CoV2 showed a significantly higher number of long disordered stretches and intrinsically disordered residues than those interacting with the other two coronaviruses, followed by the non-interacting group (Table 1). The SARS-CoV2 interacting proteins were also found to contain a higher number of long disordered protein binding regions, and residues that promote disordered protein binding than that of the other two coronaviruses, as revealed by ANCHOR2 program under IUPred2A (Table 1). However, when we compared the length of human proteins, we observed that the SARS-CoV2-interacting human proteins are significantly longer (Table 1) than those interacting with the SARS-CoV, MERS-CoV and noninteracting groups (*P*= 3.677× 10^−110^, Kruskal-Wallis test). These longer human proteins having more intrinsically disordered residues harbor disordered protein binding sites that may be utilized by the SARS-CoV2 proteins, assisting in the low-affinity protein-protein interactions with the human host.

**Table 1.** The status of protein intrinsic disorder and disordered protein binding regions of coronavirus-interacting human proteins.

| **Interaction status of human proteins** | **Mean long disordered regions** | **Average residues in long disordered regions** | **Average number of disordered residues** | **Mean long disorder-binding regions ANCHOR2** | **Average No of all disordered binding residues** | **Mean MoRF regions** | **Average number of MoRF residues** | **Protein length** |
| --- | --- | --- | --- | --- | --- | --- | --- | --- |
| **MERS-CoV interacting** | 1.123 | 99.712 | 148.651 | 2.870 | 90.192 | 0.943 | 13.486 | 566.25 |
| **SARS-CoV interacting** | 1.177 | 98.650 | 154.635 | 2.854 | 91.860 | 0.941 | 13.137 | 643.79 |
| **SARS-CoV2 interacting** | 1.626 | 139.521 | 206.838 | 3.836 | 128.163 | 1.021 | 14.433 | 725.61 |
| **SARS-CoV and SARS-CoV2 interacting** | 1.371 | 119.716 | 179.506 | 3.233 | 108.345 | 0.929 | 13.256 | 686.71 |
| **Noninteracting** | 1.017 | 84.755 | 130.925 | 2.376 | 80.017 | 0.924 | 12.585 | 523.76 |
| **P- value (Kruskal-Wallis)** | $1.356 \times 10^{-55}$ | $2.420 \times 10^{-55}$ | $1.595 \times 10^{-73}$ | $1.298 \times 10^{-53}$ | $6.298 \times 10^{-50}$ | $4.295 \times 10^{-2}$ | $4.00 \times 10^{-6}$ | $3.677 \times 10^{-110}$ |

### 3.4. Molecular Recognition Features

The protein-protein interactions involving the intrinsically disordered proteins are often mediated by disordered protein binding regions that undergo disorder-to-order transition during the physical interaction with their binding partners. These regions, better known as Molecular Recognition Features (MoRFs), contain amino acid residues that are important for protein-protein interactions (Disfani et al., 2012; Van Der Lee et al., 2014). The amino acid residues in an intrinsically disordered protein can be classified into MoRF and non-MoRF residues, with the former playing vital roles in protein-protein interaction (Panda et al., 2017), including the pathogen-host protein-protein interactions during infections (Acharya & Dutta, 2021).The abundance of MoRF residues in human-disordered proteins interacting with coronaviruses was calculated using the fMoRFpred webserver (Yan et al., 2016). Similar to our ANCHOR2 results, it was observed that the SARS-CoV2-interacting human disordered proteins contain a higher number of MoRF regions and residues, which may contribute to their interaction with viral proteins (Table 1). This also suggests that the SARS-CoV2 proteins involved in human-virus interactions utilize the disordered protein-binding MoRF regions of host proteins more extensively than those of other coronaviruses, thereby facilitating their association by promoting low-affinity interactions.

### 3.5. Evolutionary rate

The evolutionary rate of a protein depicts the change in its amino acid sequence over the evolutionary timescale. Highly expressed proteins evolve slower, due to the functional constraints imposed on them, particularly because of their high cellular demand (Drummond et al., 2005). Such proteins are more often targeted by pathogenic proteins, as their slower rate of evolution is beneficial to the pathogens for sustained interaction over evolutionary timescale. To explore further, we have studied the evolutionary rate of human proteins interacting with different coronaviruses, using 1:1 orthologs of four species: Chimpanzee, Macaque, Pig and Mouse (see materials and methods). Among all the groups, the noninteracting group was found to evolve most rapidly, and among the coronaviruses, SARS-CoV-interacting human proteins were found to evolve faster than SARS-CoV2 (Figure 2, *P*< 0.05 for all 1:1 orthologous group, Kruskal-Wallis test), suggesting that SARS-CoV2 interacts with human proteins encoded by evolutionarily more conserved genes, and therefore has a better opportunity for long-term association with its human host.

**Figure 2.**
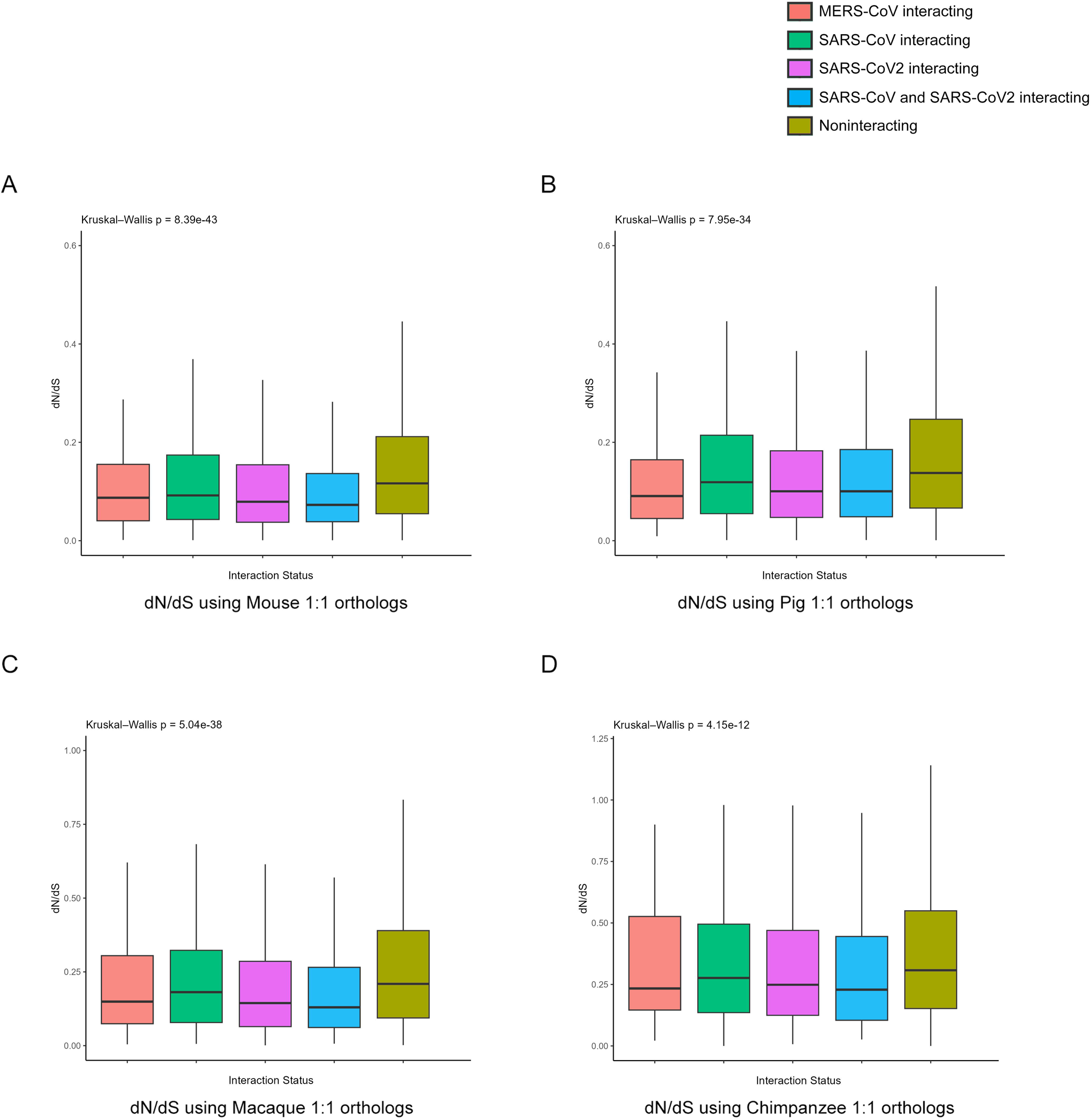
Evolutionary rate of human proteins, grouped according to the coronavirus interacting status. dN/dS values with 1:1 orthologs considering **A.** Mouse, **B.** Pig, **C.** Macaque and **D.** Chimpanzee were used to calculate the evolutionary rate of proteins.

### 3.6. Pathway enrichment analysis

During the viral infection, the host proteins that interact with the viral proteins can offer insights into the virus-utilized host pathways and functions. To explore the functional and pathway enrichments in the three coronaviruses under study, the R package clusterProfiler (Yu et al., 2012) was used to compare human proteins interacting only with one of the three coronaviruses. Enrichment analysis using the Gene Ontology Biological Process suggests that the proteins interacting with SARS-CoV2 were significantly enriched in processes like ribosome biogenesis, organelle localization, and RNA metabolism-related processes, including RNA splicing, and RNA processing, highlighting a strong association with RNA regulatory machinery. Furthermore, enrichment of chromosome segregation and spindle organization profiles suggest a possible influence of SARS-CoV2 in host cell cycle regulation (Figure 3A). In contrast, SARS-CoV showed enrichment in processes linked to chaperone-mediated protein folding and protein glycosylation, implicating modulation of host protein quality control and post-translational modification systems (Figure 3A). On the other hand, proteins interacting with MERS-CoV (MERS) were enriched in ribosome biogenesis, mitochondrial translation, and protein localization to the nucleus, indicating a functional preference for components of the host translational machinery and nuclear transport. In addition, they are also involved in viral transport and DNA structure alteration (Figure 3A).

**Figure 3.**
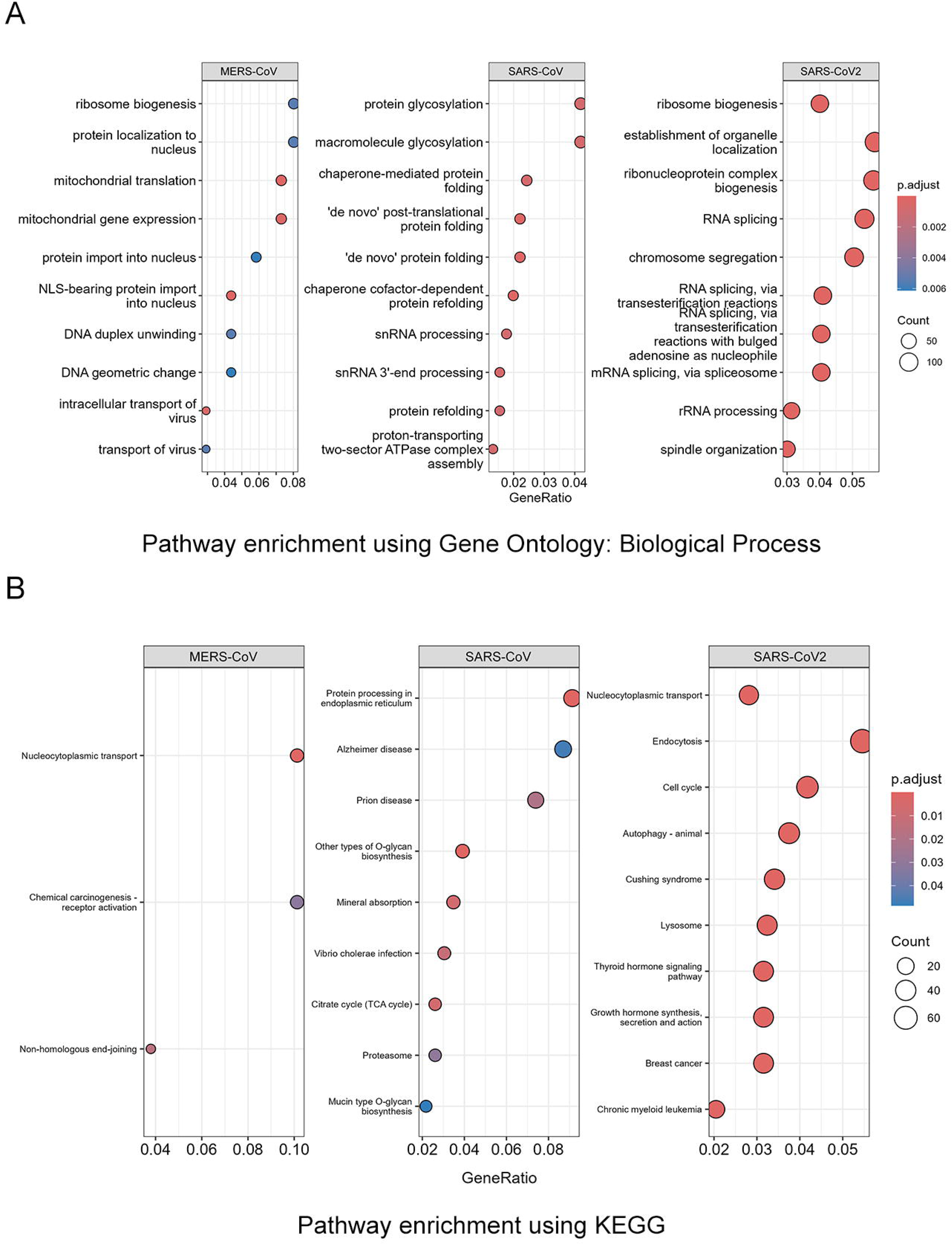
Pathway analysis of human genes interacting with the three coronaviruses: MERS-CoV, SARS-CoV and SARS-CoV2 using **A.** Gene Ontology (GO) Biological Process and **B.** Kyoto Encyclopedia of Genes and Genomes (KEGG) pathways. Dot plots display the top 10 significantly enriched pathways, with Benjamini–Hochberg adjusted P-values represented by colour gradients.

KEGG pathway enrichment further differentiated the host functional landscapes targeted by each of the viruses. The SARS-CoV2-interacting human proteins showed enrichment in nucleocytoplasmic transport, endocytosis, cell cycle and lysosome, highlighting a role in the cellular transport system (Figure 3B). In contrast, SARS-CoV-associated proteins showed enrichment in protein processing and disease-related pathways such as Alzheimer’s disease and prion disease suggesting potential overlap with host neurodegenerative mechanisms (Figure 3B). While MERS-CoV-interacting proteins were enriched in pathways including nucleocytoplasmic transport, chemical carcinogenesis and non-homologous end joining, pointing to a possible engagement with host DNA damage response and oncogenic signaling routes (Figure 3B). Together, these enrichment patterns suggest that SARS-CoV2 engages extensively with host intracellular transport and regulatory machinery, potentially facilitating efficient viral replication and immune evasion, and may reflect its broader host adaptability and fitness compared to SARS-CoV and MERS-CoV.

### 3.7. Codon usage

The function of genes relies on their encoded proteins, which fold into the desired three-dimensional structure attributed to the interactions between its constituent amino acids. However, the desired expression level of a protein is crucial for its function, often represented as the ‘translational efficiency’. After the mRNA recognition by the ribosomes, an efficient translation rate depends on the abundance of aminoacylated tRNAs, which interact with codons on mRNAs via their anticodons. Therefore, all possible codon combinations that encode the same amino acid chain do not necessarily have the same translational efficiency, though they share identical sequences (Victor et al., 2019). Viral pathogens, being dependent on host cellular machinery and the host’s aminoacylated tRNAs for translating of their proteins, should prefer a higher translation rate to facilitate their multiplication and propagation in new host cells/tissues (Nambou et al., 2022). This can be facilitated by using a similar codon usage pattern with host proteins. Here we used the EMBOSS tool to determine the frequency of each codon from MERS-CoV, SARS-CoV, and SARS-CoV2. The codon usage pattern was correlated with human housekeeping [HK] (N= 2070) and lung-specific [Lung] (N= 186) genes using the Pearson correlation coefficient (PCC) to understand the codon adaptability of these coronavirus genes. The SARS-CoV2 showed the lowest codon adaptation (PCC_HK_= 0.259; PCC_lung_= 0.052) in contrast to the MERS-CoV (PCC_HK_= 0.349; PCC_lung_= 0.172) and the SARS-CoV (PCC_HK_= 0.401; PCC_lung_= 0.213) with both the housekeeping (*P* < 0.05 for PCC), but not the lung-specific gene groups (*P* > 0.05 for PCC) (Figure 4). This suggests that both the SARS-CoV and MERS-CoV show better codon adaptation with HK genes but show an insignificant trend with Lung genes, compared to the SARS-CoV2.

**Figure 4.**
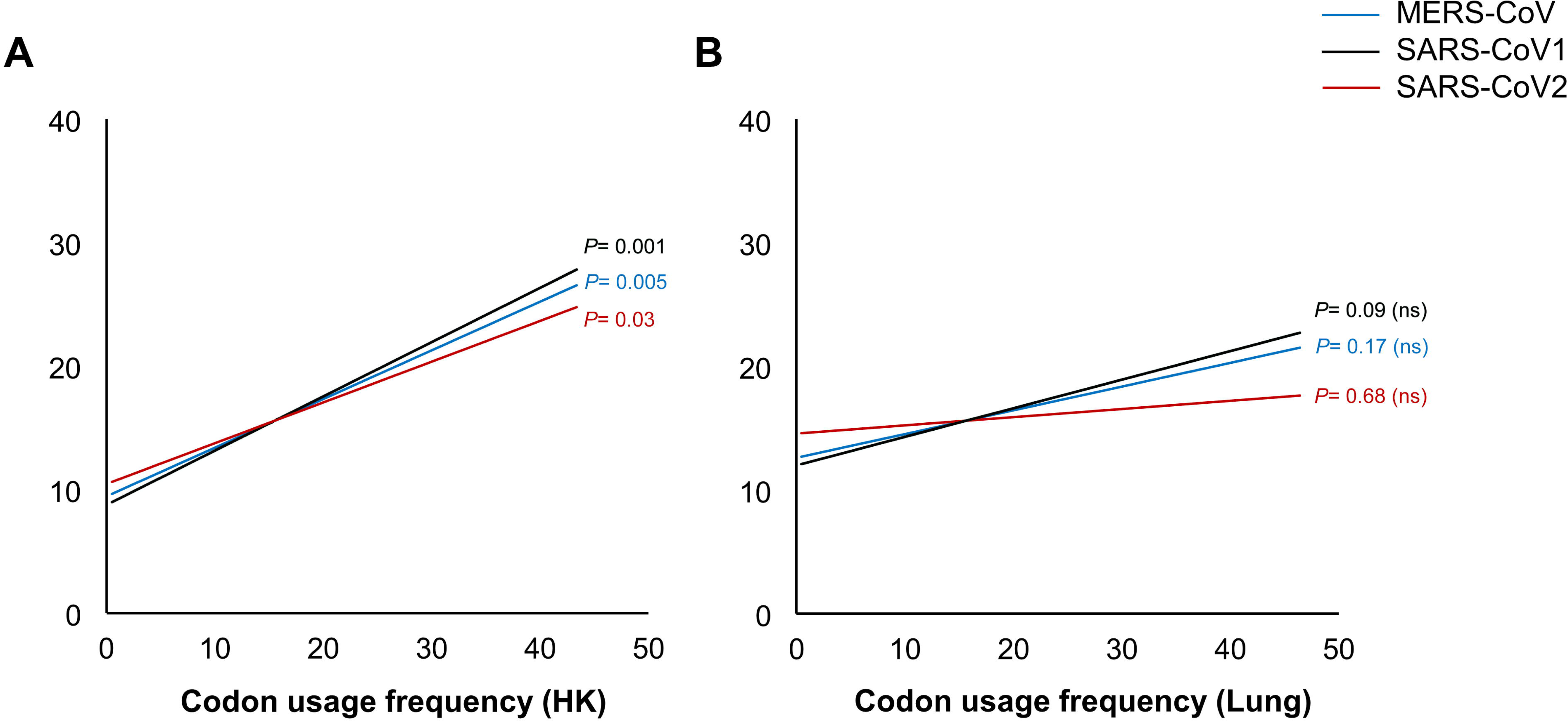
Codon usage correlation of MERS-CoV, SARS-CoV and SARS-CoV2 with the human genes **A.** housekeeping genes and **B.** lung-specific genes, using Pearson correlation coefficient. MERS-CoV is represented in blue while SARS CoV in black and SARS-CoV2 in red.

Although SARS-CoV2 exhibits widespread interaction with human proteins, its lower codon adaptation to human housekeeping and lung-specific genes suggests that it has not yet undergone extensive evolutionary optimization for translational efficiency in the human host. This discrepancy may stem from the virus’s recent zoonotic spillover, giving it less evolutionary time to fine-tune its codon usage in alignment with host tRNA pools. In contrast, SARS-CoV and MERS-CoV, which have either circulated longer or undergone more cycles of host adaptation, demonstrate higher codon usage correlation with human genes, potentially facilitating more efficient translation.

## 4. Discussion

The host-pathogen coevolution has been a popular area of research for ages, where the hosts are exploited by the pathogens for their survival and reproduction, causing several ailments including life-threatening diseases. Although host-pathogen interaction is better explainable under their coevolutionary dynamics, it is difficult to portray due to the complex nature of the interactions at the cellular, molecular, and biochemical levels, and the availability of only a snapshot of these interactions at any point in time. Of particular importance are viruses that utilize their host cellular machinery to synthesize the proteins required for survival, reproduction, immune evasion, and disease progression (Walsh & Mohr, 2011). For a deadly virus like SARS-CoV2, it is not easy to establish a sustainable coevolutionary relationship with the novel host, as its deadly effects may kill the host, hampering the propagation of the virus. However, with time, molecular interactions between host and virus usually get stabilized, forming a stable pathogen-host protein-protein interaction (PHPPI) network, resulting in coexistence. In this study, we compared the virus-host interactome of the novel coronavirus SARS-CoV2 and the other two human coronaviruses, namely SARS-CoV and MERS-CoV from the perspective of the PHPPI network. The central proteins were previously shown to be the prime targets of pathogens and are involved in the PHPPI network (Acharya & Dutta, 2021; Crua Asensio et al., 2017; Dyer et al., 2008). These proteins play important roles in the host cellular processes and assist pathogens to gain control over the host protein-protein interaction (PPI) network. The higher infectivity of SARS-CoV2 may have shaped their host-adaptation better than the closely related SARS-CoV. Here we observed that the SARS-CoV2 interacts with human proteins that are more central, containing a higher proportion of network hubs and bottlenecks than those interacting with other coronaviruses, namely MERS-CoV and SARS-CoV, which cause milder infections. By interacting with central proteins in the host PPI network, SARS-CoV2 increases its evolutionary fitness by gaining control over host cellular systems. Delving further, we observed that the human proteins interacting with SARS-CoV2 are associated with a higher number of protein complexes, allowing them to interact with additional functional modules in the human PPI network. However, previous studies suggested that pathogens could utilize intrinsically disordered proteins or intrinsically disordered residues to facilitate low-affinity interactions (Acharya & Dutta, 2021; Blundell et al., 2020). Therefore, we explored the presence of disordered stretches present within human proteins interacting with different coronaviruses. The PHPPI network revealed a higher involvement of intrinsically disordered proteins in the SARS-CoV2-interacting human proteins, with a higher number of disordered protein binding sites promoting such low-affinity interactions, compared to the other two coronaviruses (Table 1). Moreover, we also observed that the protein length is also a major determinant of host-virus interaction, as longer proteins have a probability of harboring more intrinsically disordered residues and disordered protein binding motifs (Table 1). So, their abundance, but not density, is the key determinant, and their presence may facilitate the host-pathogen interactions. Considering the protein evolutionary rates, human proteins interacting with SARS-CoV2 are more evolutionarily conserved than the two other coronaviruses, suggesting that its PHPPI network will favour a long-term host association (Brito & Pinney, 2017).

In addition, to survive and multiply in the host’s body, the viral pathogens must utilize the host’s cellular machinery. The comparative pathway-level analysis using three coronaviruses indicates that SARS-CoV2-interacting human proteins are associated with crucial biological pathways in humans, including ribosome biogenesis, organelle localization, RNA metabolism-related processes, chromosome segregation, spindle organization, cell cycle and nucleocytoplasmic transport that are central to host cellular functions. Furthermore, another crucial component of viral adaptation to the host comes from the host codon usage preferences (Bahir et al., 2009; Nambou et al., 2022). The host tRNA pool and codon usage remain optimized for the desired expression of the host proteins (Victor et al., 2020). For a novel host-virus association, this is merely a chance factor that the host tRNA pool will support the codons used by the viral proteome. However, with time, viruses may adapt to host codon usage preferences to maintain optimum efficiency in their multiplication (Bahir et al., 2009). We correlated the coronavirus codon usage pattern with that of the human housekeeping genes and the genes enriched in the lung tissues for each of the coronaviruses. The housekeeping genes need a steady expression in all tissues and the required cognate tRNAs should be maintained at a steady level for their expression. Whereas, genes expressed in the lung, one of the sensitive sites for colonization and infection of the coronaviruses, should maintain a steady cognate tRNA pool for the optimized expression of lung-specific genes. A lower correlation (PCC) in the codon usage pattern of SARS-CoV2 with the human housekeeping genes was observed, followed by MERS-CoV and SARS-CoV. Moreover, SARS-CoV2’s broad tissue tropism and rapid transmissibility may be driven by mechanisms beyond codon optimization, such as its ability to modulate host metabolism and immune evasion pathways to compensate for suboptimal translational compatibility.

However, this study has several inherent limitations that should be acknowledged. First, the protein-protein interaction (PPI) data used in this study are representative static snapshots from heterogeneous sources, limiting the dynamic representation of host-pathogen coevolution. The interactome data are also disproportionately enriched for SARS-CoV2 due to its recent global impact and a huge number of studies in comparison to the limited datasets for SARS-CoV and MERS-CoV, potentially biasing comparative analyses. Furthermore, this study does not include strains/variants in the viral PPI datasets or temporal PPI dynamics for each virus due to the lack of specific information. Nevertheless, within the circumstantial limitations, this study highlights the importance of human-coronavirus protein-protein interaction and the role of protein structural features that can assist SARS-CoV2 with greater transmission and host-cellular control, despite sharing close genomic sequence identity with the less-virulent SARS-CoV.

## 5. Conclusion

This comparative analysis of host-pathogen protein-protein interactions (PPIs) among three major human coronaviruses reveals critical insights into viral strategies for host exploitation and adaptation. Our findings demonstrate that SARS-CoV2 preferentially targets central and functionally versatile human proteins, engages with a higher number of protein complexes, and utilizes intrinsically disordered regions and longer host proteins to form dynamic, low-affinity interactions. These characteristics enhance its capacity to interface with multiple host pathways and functional modules. Despite showing the lowest codon adaptation to the host’s translational machinery among the three viruses, SARS-CoV2 compensates by targeting essential host pathways, including those involved in RNA processing, cell cycle control, and nucleocytoplasmic transport—thereby optimizing its replication and propagation. This contrast between structural interaction patterns and translational compatibility reflects a unique adaptation strategy that may support the high transmissibility of SARS-CoV2, compared to SARS-CoV and MERS-CoV.

## CRediT authorship statement

Debarun Acharya (Conceptualization, Investigation, Formal analysis, Writing-original draft), and Tapan K. Dutta (Conceptualization, Supervision, Writing-review & editing)

## Ethical approval

This article does not contain any studies with human participants or animals performed by any of the authors.

## Data availability

Data will be made available on request.

## Acknowledgement

The authors thank the Director, Bose Institute, Kolkata, India for encouragement and support.

## Abbreviations

CDS: coding sequence
CoV: coronavirus
GO: gene ontology
HK: housekeeping
IBV: infectious bronchitis virus
MERS: Middle East respiratory syndrome
MHV: mouse hepatitis virus
MoRF: molecular recognition feature
PCC: Pearson correlation coefficient
PHPPI: pathogen-host protein-protein interaction
PPI: protein-protein interaction
SARS: severe acute respiratory syndrome
tRNA: transfer-ribonucleic acid

